# Screening for functional regulatory variants in open chromatin using GenIE-ATAC

**DOI:** 10.1101/2022.02.09.479775

**Authors:** Sarah Cooper, Jeremy Schwartzentruber, Eve L Coomber, Qianxin Wu, Andrew Bassett

## Abstract

Understanding the effects of genetic variation in gene regulatory elements is crucial to interpreting genome function. This is particularly pertinent for the hundreds of thousands of disease-associated variants identified by GWAS, which frequently sit within gene regulatory elements but whose functional effects are often unknown. Current methods are limited in their scalability and ability to assay regulatory variants in their endogenous context, independently of other tightly linked variants. Here we present a new medium-throughput screening system: <u>g</u>enome <u>en</u>gineering based <u>i</u>nterrogation of <u>e</u>nhancers <u>a</u>ssay for <u>t</u>ransposase <u>a</u>ccessible <u>c</u>hromatin (GenIE-ATAC), that measures the effect of individual variants on chromatin accessibility in their endogenous genomic and chromatin context. We employ this assay to screen for the effects of regulatory variants in human induced pluripotent stem cells, validating a subset of causal variants, and extend our software package (rgenie) to analyse these new data. We demonstrate that this methodology can be used to understand the impact of defined deletions and point mutations within transcription factor binding sites. We thus establish GenIE-ATAC as a method to screen for the effect of gene regulatory element variation, allowing identification and prioritisation of causal variants from GWAS for functional follow-up and understanding the mechanisms of regulatory element function.

## Introduction

Genome-wide association studies (GWAS) of complex diseases and traits have identified an ever-growing list of associated genetic loci currently numbering 326,947 (1). However, it remains hugely challenging to identify specific causal variants and genes, since the majority of these loci lie in non-coding regions of the genome and there are often many candidate variants in linkage disequilibrium (LD). The identification of causal genes has been aided by expression quantitative trait loci (eQTL) datasets from relevant cell types, since colocalization between eQTL and GWAS results can indicate a shared genetic cause of disease and gene expression change (2, 3). Whilst useful, these datasets are currently underpowered, resulting in many disease associated loci lacking an eQTL association to a specific gene, and in other cases, multiple genes being implicated at the same locus (4). Another method to link a disease association to a gene is through annotation of enhancer-promoter loops using chromatin conformation capture (3C, HiC, pchiC, captureC) methodologies (5), but these have limitations in resolution, frequently missing short-range interactions (6), and often implicating multiple genes (7).

Establishing causal variants is perhaps even more challenging. Statistical fine-mapping methods can be used to narrow down the list of putative causative SNPs from human genetic data (4, 8), but their resolution is limited by the recombination structure in human populations. Sites of open chromatin, marked by accessibility to DNA modifying enzymes (e.g. DNAseI hypersensitivity (9) or assay for transposase accessible chromatin, ATAC (10)), are bound by transcription factors (TF) in place of canonical nucleosomes. Variants residing at these genomic locations could therefore exert their effects by altered TF binding and gene regulation. Thus, annotation of chromatin features, such as open chromatin domains and histone modifications, can also help to identify likely causative SNPs and provide an insight into the molecular mechanism (11). However, such chromatin features are highly cell type and stimulation specific, making it difficult to rule out particular variants, and frequently implicate multiple candidate variants. Chromatin QTL studies are able to identify regulatory DNA variation that directly impacts chromatin architecture (12) but similarly to eQTL analysis, these studies are often underpowered and do not necessarily implicate individual variants (6). Importantly, none of these methods are able to directly demonstrate the causality of a specific variant. Massively parallel reporter assays can be used to understand the effect of specific variants on enhancer activity (13, 14), but enhancers are removed from their native genomic and chromatin context, and thus are unable to demonstrate that a specific variant truly has an effect *in vivo*. Use of genome engineering technologies to introduce specific SNPs is an attractive way to causally link a variant to a change in enhancer or gene function (15), but these experiments are time-consuming and difficult to scale. It is also possible to use CRISPR inhibition or activation techniques targeted to presumed enhancer regions to screen for those that are active in a particular cell type (16–18), but the resolution of these methods is limited to around 1 kb, and they do not show an effect of a specific genetic variant(s).

To circumvent some of these limitations, we have previously developed a medium throughput arrayed screening method we termed genome engineering based interrogation of enhancers (GenIE) that allows screening for the effect of a variant of interest on transcription of the gene in which it resides (19). However, this is limited to those variants present within transcribed regions of the genome, which does not allow analysis of the large number of intergenic enhancers, in which GWAS variants are selectively enriched (20). Here we have developed a method, GenIE-ATAC, that allows screening for the effect of specific genetic variants on the open chromatin region in which they are located, in their endogenous context.

## Results and discussion

We extended our existing GenIE method (19) to assay the effect of variants lying within gene regulatory elements on the chromatin accessibility of that region. We introduced a variant of interest by genome editing, and used a locus-specific ATAC to measure the effect of each allele on chromatin accessibility. Thus, we could directly demonstrate the causality of a specific variant in altering chromatin at its endogenous location within the genome. This allowed us for instance to distinguish between several causal variants in high LD, or to establish cause and effect relationships for co-regulated enhancers. In contrast to the alternative approach of making edited isogenic cell lines, we were able to avoid the time-consuming step of clonal isolation (21) and the problem of clone-to-clone variability (22) by analysing the pool of edited cells. This made it possible to perform an arrayed screen of dozens of variants, and will, in the future, allow the use of more complex and disease relevant cellular models such as differentiated induced pluripotent stem cells (iPSCs) and primary cell types.

We used human iPSCs to establish our method, since these cells can be used to model regulatory element variation in a wide variety of disease-relevant cell types and are amenable to CRISPR-mediated genome editing. We first delivered Cas9 and a guide RNA as a ribonucleoprotein along with a 100 nt single-stranded DNA oligonucleotide homology directed repair (HDR) template (21). This generated a population of cells containing the WT allele (unedited) and the HDR allele (SNP introduced) along with a large number of indels (small insertions/deletions) (Fig 1).

**Figure 1.**
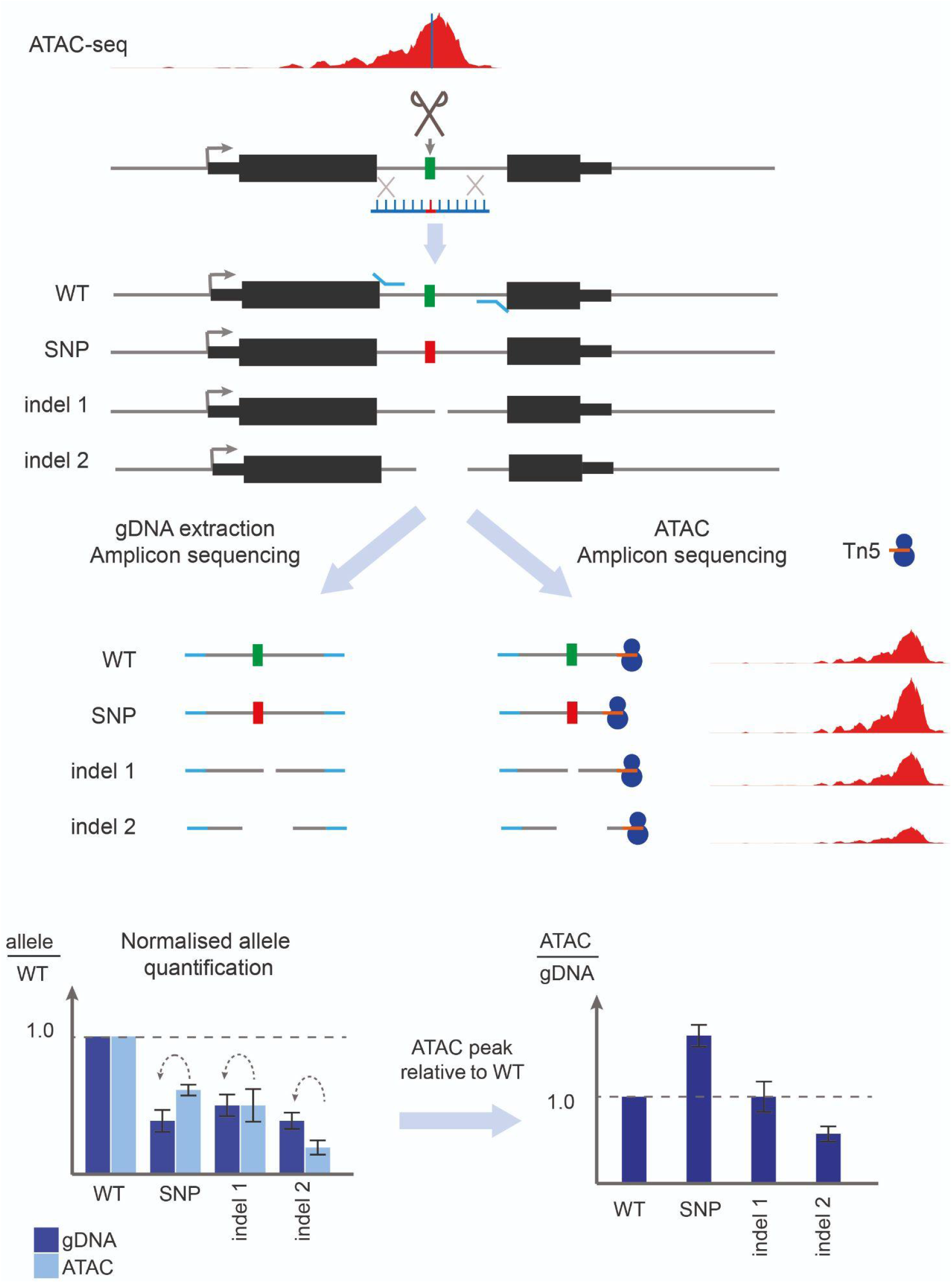
Schematic of GenIE-ATAC assay. Edited pools of cells contain a mixture of WT, edited point mutation (SNP) and a variety of deletion alleles (indel 1, 2 etc.). The chromatin accessibility (and therefore transcription factor occupancy) of each of the genotypes can be quantified by amplicon sequencing of the ATAC material and gDNA extracted from the same population of cells. Values are internally normalised to the WT allele for each library and displayed as a ratio of ATAC to gDNA reads as a measure of relative chromatin accessibility for each allele.

We harvested genomic DNA (gDNA) from the mixed population of cells, performed PCR across the edited site and assessed the frequency of different alleles within the population using high throughput sequencing of the amplicon. Using exactly the same population of cells we performed ATAC, which selectively integrates adaptor sequences into open chromatin using the Tn5 transposase. We then carried out a nested PCR using a target-specific primer near to the edited site and a universal primer that binds the Tn5 adaptor sequence. This amplicon was also sequenced, and the effect on chromatin accessibility for the edited allele was then calculated as the ratio of sequencing reads in the ATAC relative to the gDNA for the edited allele, normalised to the same ratio for the unedited (WT) allele.

Two developments were necessary to successfully establish a site-specific ATAC protocol. Firstly, we purified our own Tn5 and loaded it with a single oligonucleotide sequence (MEDS-A sequence) to allow a single PCR primer binding site for the subsequent PCR of the ATAC sample. Our method also requires a substantial input of ATAC DNA into the PCR to avoid any PCR bias, and the cost of commercial Tn5 protein would limit the number of variants that could be assayed in a single experiment. Secondly, we needed to carry out a linear PCR using a biotinylated primer, followed by a biotin pulldown of the amplified DNA to enrich for the region of interest prior to a semi-nested PCR (Fig S1). This is because the Tn5 MEDS-A sequence is integrated at tens or hundreds of thousands of sites in the genome, whereas the gene specific primers only have one binding site, and thus a semi-nested PCR alone was unable to give sufficient enrichment of the desired region.

We first tested GenIE-ATAC on three heterozygous SNPs in KOLF_2 iPSCs that overlapped with accessible chromatin regions defined by ATAC-seq in these cells. These variants were selected due to their high probability of affecting chromatin accessibility based on fine-mapping of chromatin QTL data from sensory neurons (23). As expected for heterozygous alleles, when we performed amplicon sequencing over the variant of interest from gDNA we observed an equal proportion of the two alleles (ratio of 1) (Fig 2, red bars).

**Figure 2.**
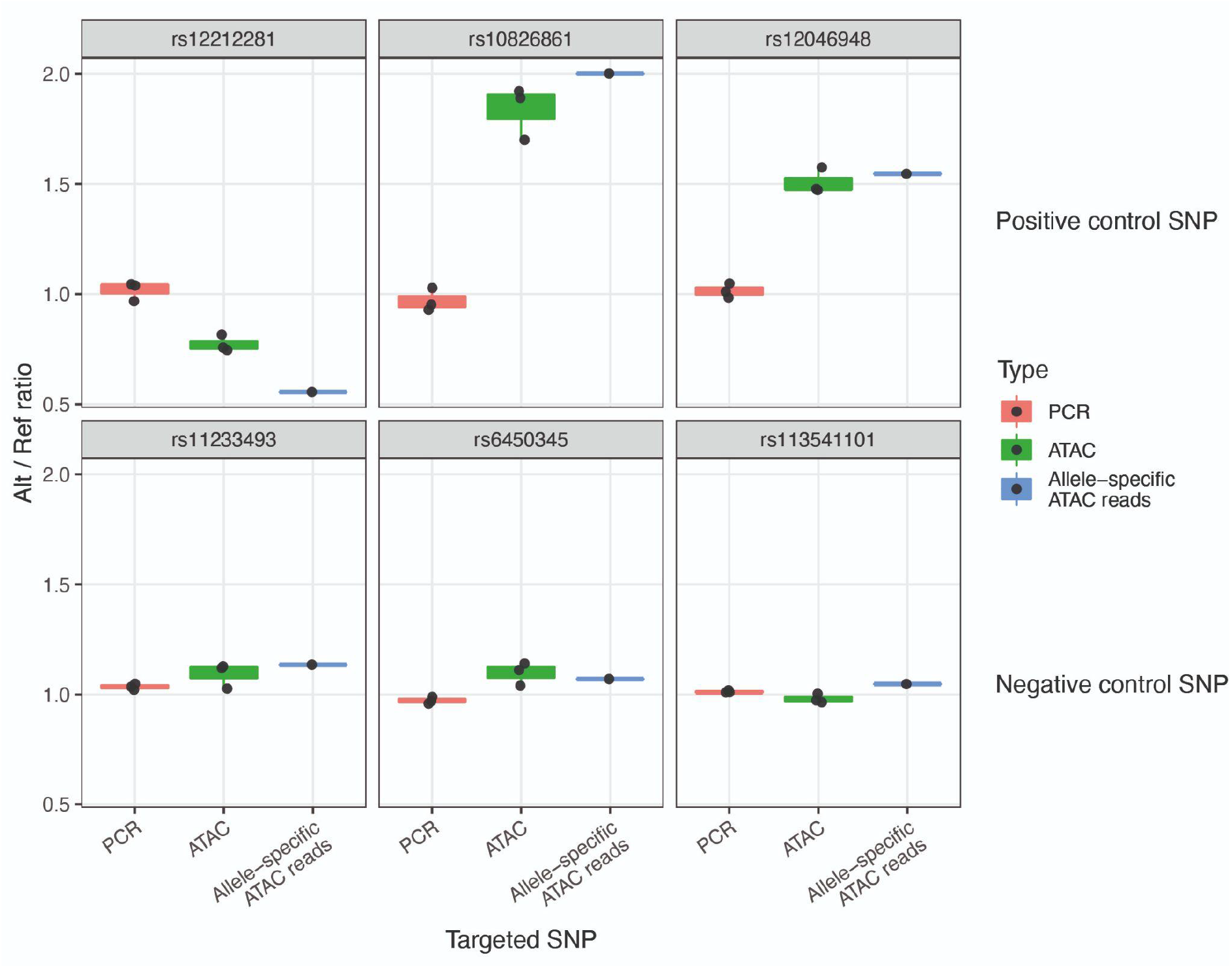
GenIE-ATAC recapitulates allele specific SNP effects on chromatin accessibility. GenIE-ATAC was performed on KOLF_2 iPSCs at six heterozygous SNPs: three positive control SNPs (top) with predicted effects on chromatin accessibility based on chromatin QTL analysis in iPSCs, and three negative control SNPs (bottom) that would not be predicted to have an effect. Graphs show the ratio of chromatin accessibility of the alt allele normalised to the reference genome sequence (Alt/Ref ratio). Red bars indicate amplicon sequencing of gDNA (PCR), green bars from amplicon sequencing of ATAC material (ATAC), blue bars show allele-specific reads from ATAC-seq data (allele-specific ATAC reads) from KOLF_2 iPSCs. Individual repeats are indicated with dots; boxplot hinges represent the 25th and 75th percentiles, where these are interpolated due to the small number of points (n=3), and whiskers extend to the most extreme data point not further than 1.5 times the inter-quartile range from the hinge.

For each SNP tested, when we analysed amplicon sequencing using GenIE-ATAC, there was an imbalance of reads between the two alleles (ratio skewed away from 1), with the alt alleles either decreasing chromatin accessibility (for SNP1) or increasing it (for SNPs 2 and 3). These results were recapitulated using primers designed in the reverse direction with the locus specific primer on the opposite side of the variant (Fig S2). Importantly, the effects of SNPs on chromatin accessibility measured by GenIE-ATAC were consistent with the expectations from the chromatin QTL data and confirmed independently by analysis of allele-specific reads from ATAC-seq in hiPSCs (19). As negative controls for GenIE-ATAC we also performed the experiment using three heterozygous SNPs located in ATAC peaks that had no strong allelic imbalance in ATAC-seq (Fig 2, bottom panel), and once again the data from GenIE-ATAC corroborated that seen with allele-specific ATAC-seq.

The control in these experiments was a simple PCR from gDNA, but we wanted to validate that we obtained the same results with tagmented gDNA that had been through the linear PCR and enrichment process. We found that tagmentation and enrichment did not cause an allelic bias, and we still observed a ratio of 1 for heterozygous alleles. However it did increase the variation we observed (Fig S3) when compared to simple gDNA PCR, likely due to sampling bias. Thus, we used gDNA PCR as the control for subsequent experiments to increase consistency and the throughput of the assay. In order to increase the number of replicates that could be achieved per SNP per experiment, we also tested the effect of titrating the amount of ATAC DNA to be added into the linear PCR. The results and variation were similar with 3 times less input material (Fig S4), which we used for subsequent experiments.

We next used GenIE-ATAC to screen 8 naturally occurring SNPs for their causal effect on accessibility of chromatin after editing with CRISPR. These SNPs were selected based on being homozygous, located in ATAC peaks in our iPSC line, and having a high probability of affecting chromatin accessibility from chromatin QTL data (23). The details of these regions and their predicted effects on transcription factor binding (24) are indicated in Table 1.

**Table 1.** Details of edited SNPs and predicted effect on transcription factor binding.

|  | SNP ID | Position of SNP | Nearest gene | Position of SNP with iPS ATAC peak | KOLF2 genotype | Edit introduced by HDR | Binding site information |
| --- | --- | --- | --- | --- | --- | --- | --- |
| 1 | rs55685444 | chr5:33470746 | TARS | 0.54 | C hom ref | G alt | In motifs of CTCF and JUND, within binding peaks. DeepSEA predicts decrease in CTCF binding by the alt allele. |
| 2 | rs12269414 | chr10:63872514 | REEP3 (250 kb away) | 0.61 | G hom ref | A alt | In a CTCF motif within a CTCF peak. DeepSEA predicts massively decreased CTCF binding for the alt allele. |
| 3 | rs7729529 | chr5:43007716 | ZNF131 and SEP1 | 0.17 | C hom ref | G alt | In a region of dense TF binding. DeepSEA predicts significantly increased binding of CEBPB. |
| 4 | rs6700034 | chr1:20508117 | MUL1 | 0.33 | C hom ref | A alt | In 5'UTR of MUL1. Alt allele had a transcriptional increase in GenIE assay. |
| 5 | rs8011143 | chr14:22599022 | ABHD4 | 0.52 | T hom ref | C alt | Within intron 1 of ABHD4. Alt allele had a transcriptional decrease in GenIE assay. |
| 6 | rs4757506 | chr11:17276730 | NUCB2 | 0.49 | A hom ref | G alt | SNP very near the start of the 5' UTR. DeepSEA predicts slight decreases in CTCF binding and histone marks. |
| 7 | rs185220 | chr5:56909530 | SETD9 | 0.26 | G hom alt | A ref | In a region of dense TF binding in the 5' UTR of SETD9. DeepSEA predicts strongly increased DNaseHS for the alt allele, as well as multiple TFs. |
| 8 | rs141252451 | chr11:11799003 | MIR8070 | 0.51 | hom ref | Insertion CA alt | SNP is right in the small peak centre, in a CTCF motif and Chip-seq peak. DeepSEA predicts completely lost CTCF binding. |

The editing efficiency of each of these SNPs in our WT iPSC line (KOLF_2) was variable but generally high (Fig 3A, bottom panel). This may be because they are located in regions of accessible chromatin that have previously been shown to have particularly high editing rates (25). This is advantageous for GenIE-ATAC since all variants will be located within ATAC peaks. Data was processed using an R package we have developed (rgenie) which automates the analysis of experiments, and reports quality control and statistical analysis of the effect of both HDR alleles and defined small deletions over the variant of interest.

**Figure 3.**
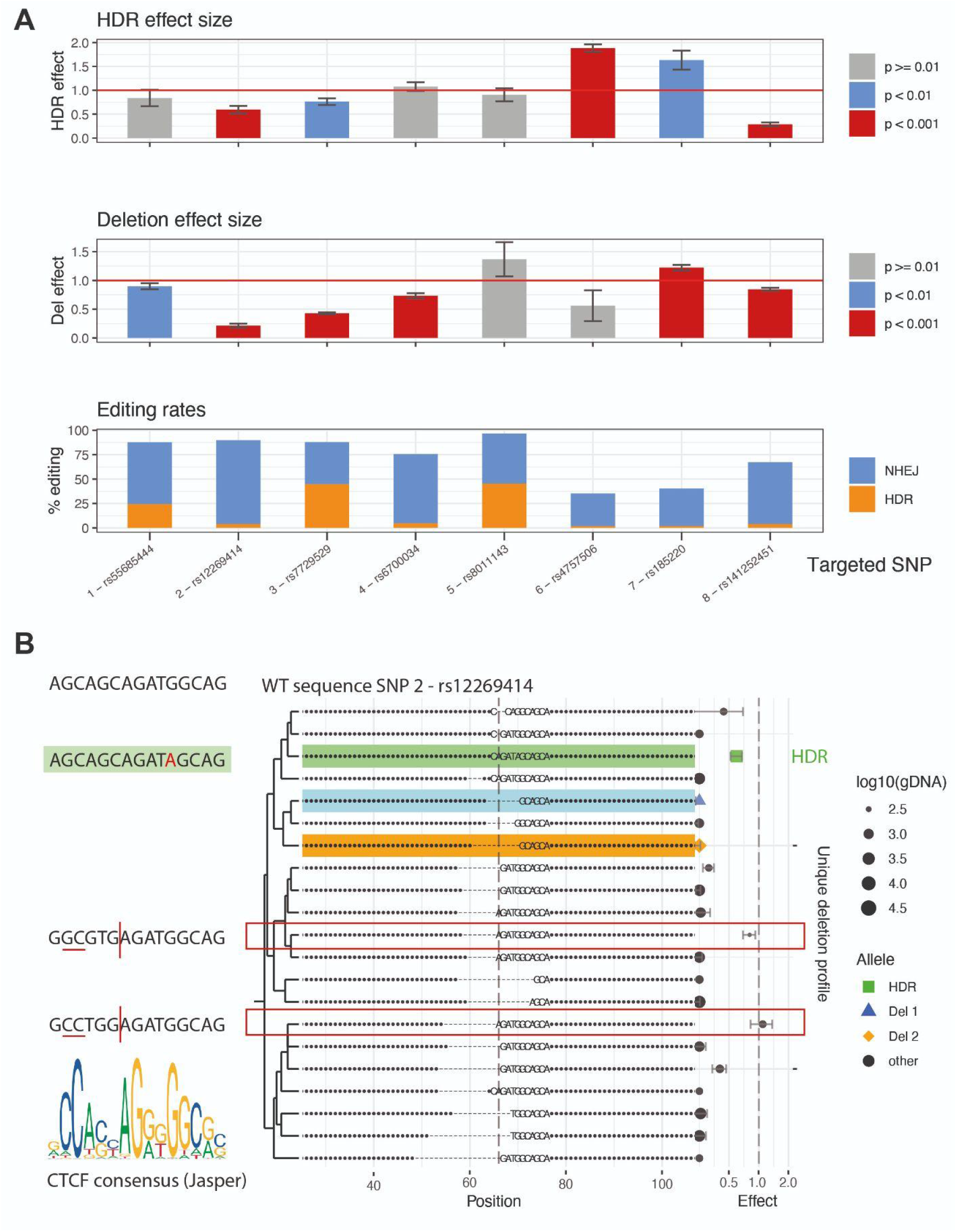
GenIE-ATAC CRISPR screening identifies functional regulatory SNPs. **A.** Homozygous SNPs (1-8) were edited using CRISPR to their alternative alleles in KOLF_2 hiPSCs and assayed by GenIE-ATAC. The top panel shows the effect on chromatin accessibility of the HDR-introduced allele relative to the WT allele. The middle panel shows the effect size of deletions within a +/− 6 bp window of the SNP site relative to the WT allele. The bottom panel shows editing rates (HDR, orange; indels, blue). Data was processed using rgenie. All error bars represent 95% confidence intervals, n=3. **B.** The most frequent deletion types by read count from GenIE-ATAC targeting of SNP 2 (rs12269414). The effect size of ATAC for each allele relative to the WT is indicated on the right hand side. The HDR allele is highlighted in green. Del 1 and Del 2 (highlighted in blue and orange respectively) are small defined deletions around SNP of interest. Deletion alleles which have no effect change are indicated by red boxes, and recapitulate the CTCF motif (left).

The HDR effect size after editing the desired SNPs to the alternative allele (Fig 3A, upper panel, and Supplementary Table 3) showed that three of the edited SNPs did not have any significant effect on chromatin accessibility (Regions 1, 4 and 5). We have previously shown that the SNP in region 4 has an effect on transcription of the MUL1 gene (19), so a lack of change in chromatin accessibility suggests its effect is mediated post-transcriptionally, likely at the RNA level. SNPs edited in regions 2, 3 and 8 all showed a decrease in chromatin accessibility consistent with a reduction in transcription factor binding. The Region 2 SNP is within a CTCF binding motif and the editing of a G>A within the motif is consistent with a predicted reduction of CTCF binding and chromatin accessibility upon editing. Region 8 had the largest effect size in the screen, and in this case the HDR event created a 2 bp insertion (CA) within a CTCF binding site, which was predicted to result in the complete loss of CTCF binding. Interestingly, the SNPs in regions 6 and 7 showed an increase in chromatin accessibility, potentially due to the introduction of a new binding site for a transcription factor.

As well as introducing the SNP of interest at each site by HDR, the GenIE-ATAC method also creates a large population of deletion and insertion alleles around the cut site of the Cas9/guide. We can use these to provide information about the effect of these mutations on chromatin accessibility. Accordingly, we analysed the effect of all deletions wholly contained within a 6 bp window around the SNP of interest (Fig 3A, middle panel, and Supplementary Table 3). As expected, deletion of bases within an ATAC peak very often results in a reduction of chromatin accessibility due to disruption of transcription factor binding, and we observed this in 6 of our 8 edited locations. However, deletions around the SNP in region 5 did not have any effect and deletions in region 6 had an opposite effect to the introduction of the SNP, demonstrating the importance of being able to introduce and assay specific desired variants. Closer inspection of individual deletions in region 4 (Fig S5) showed that those that removed a poly-C region had a strong effect on chromatin accessibility, but those that did not had a weak or no effect. This suggests that these C bases are important for binding of an unknown factor, and highlights how analysis of specific deletions can provide information about mechanism of action. Analysis of region 2 (Fig 3B), demonstrated that whilst introduction of the SNP led to a loss of accessibility, the deletions fell into two classes where accessibility was virtually abrogated, or was retained (red boxes). Closer inspection showed that in the case of the specific deletions which maintained chromatin accessibility, the adjacent sequences were such that the CTCF consensus site was recapitulated upon deletion. Thus, it is highly likely that CTCF binding was responsible for the accessible chromatin peak at this site. All plots were generated using the rgenie R package and the full output for all SNPs in this experiment is shown in supplementary note rgenie_output.

The effects of point mutations within transcription factor binding sites are often difficult to predict based on consensus sequences alone (26). We therefore performed GenIE-ATAC in a multiplex format to simultaneously introduce multiple SNPs across a defined region in one experiment and obtain nucleotide-level resolution of the determinants of transcription factor binding. We targeted the CTCF binding site in region 2 and designed HDR oligo templates to mutate 9 bases in the binding site individually into every other possible base (Fig 4A).

**Figure 4.**
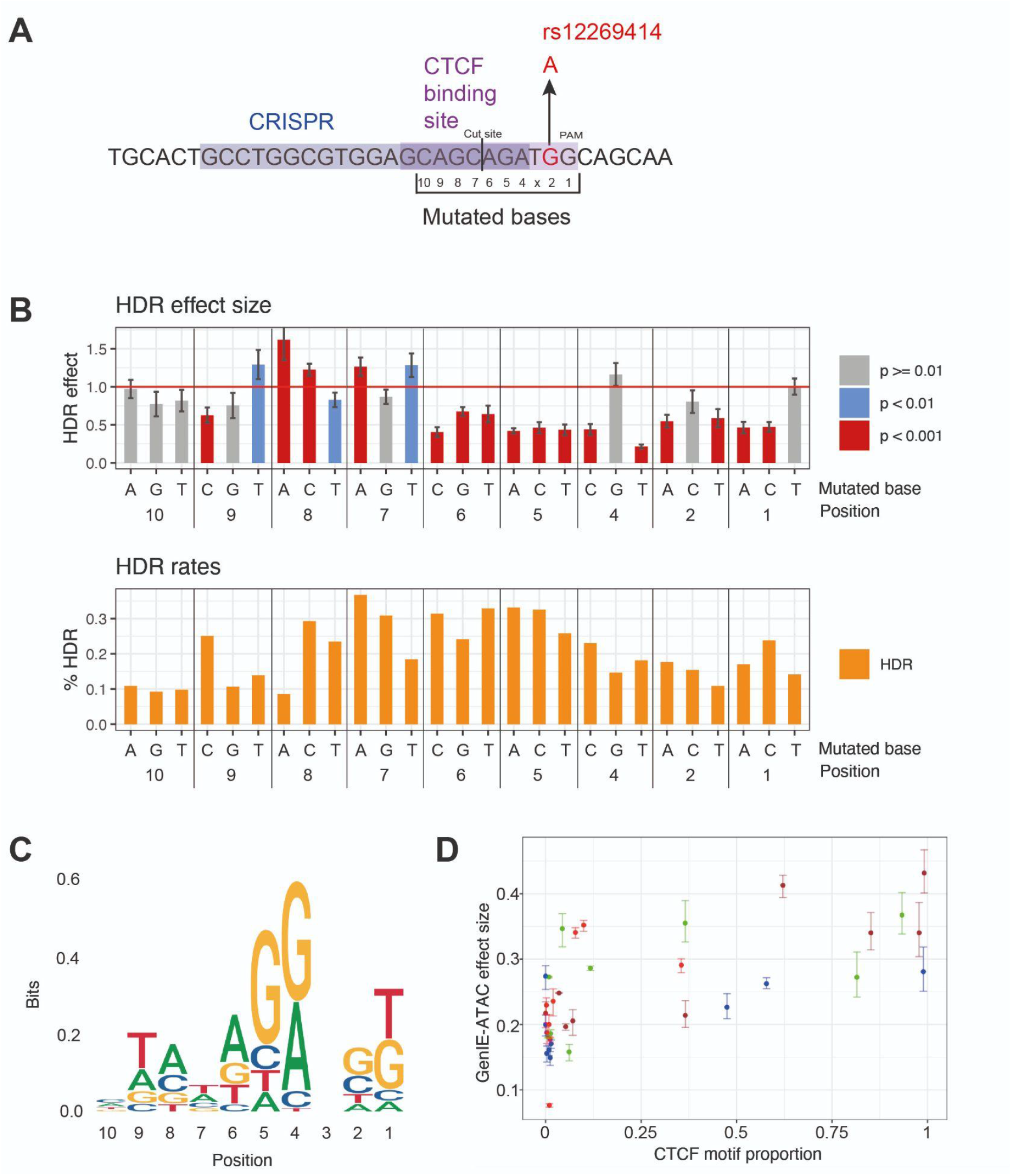
GenIE-ATAC across a CTCF binding site highlights SNPs important for binding. **A.** Schematic of the CTCF binding site around rs12269414 and mutated bases. Position 3 is the N within the NGG PAM sequence of the CRISPR and therefore could not be mutated due to recutting and subsequent indel formation. **B.** Top panel shows the effect on chromatin accessibility of the HDR allele normalised to the WT allele with mutated bases indicated at the bottom. Bottom panel shows editing rates. Data was processed using rgenie. All error bars represent 95% confidence intervals, and p values are as indicated. **C.** Sequence logo showing the consensus binding site as determined by GenIE-ATAC, with letter size proportional to the square of the relative GenIE-ATAC effect size when mutated to that nucleotide (relative to the WT/consensus, set to 1). **D.** Scatter plot showing correlation between GenIE-ATAC measured effect size and Jasper CTCF motif proportion size. Points represent the GenIE effect size for each nucleotide divided by the sum over all 4 nucleotides at the same position; error bars are the 95% confidence intervals for a given nucleotide divided by the sum of these over all 4 nucleotides at the same position.

We performed GenIE-ATAC using a pool of all HDR oligos. It was not possible to introduce mutations at the N in the NGG PAM due to the fact that Cas9 would continue to introduce indels even after successful HDR, and bases further from the Cas9 cut site could not be evaluated due to low HDR rates. We increased the amount of ATAC DNA material and gDNA into each PCR proportionally for the number of HDR events to maintain representation (see methods). Fig 4B shows the effect size of each substitution (see also Supplementary Table 4). For example, a substitution of an A to G in position 4 of the CTCF site had no effect on binding, whereas mutation to any other base dramatically decreased chromatin accessibility. The inverse of this effect size reflects the importance of each base for CTCF binding, and broadly recapitulates the CTCF binding motif (Fig 4C, D). There are some differences, however, which could be due to several reasons. Firstly, the consensus binding motif measures the frequency of each base across all binding sites, and the nucleotide determinants of binding at a specific locus may be different (e.g. due to competitive or cooperative binding interactions, effects of DNA major/minor groove structure, or chromatin modifications or structure). Secondly, DNA binding factors other than CTCF could be contributing to the changes in chromatin accessibility. Finally, the effect of mutation has been shown to be highly context-specific, such that mutation of a consensus base can be rescued or buffered by a second mutation elsewhere in the binding site (26). This result illustrates the context-dependence of transcription factor binding and the difficulty in predicting the effect of a mutation from binding site consensus alone.

## Conclusion

We believe that GenIE-ATAC will be a generally useful screening method to identify causal variants from GWAS studies and to understand transcription factor binding in the context of chromatin. Importantly, it is able to functionally link specific genetic variants to changes in chromatin accessibility in their endogenous locus and genomic context. It can thus validate causal variants, and provide mechanistic information about how they could influence transcription factor binding and gene expression, which is currently understood in a vanishingly small number of cases. Advances in genome editing methods such as improved homology directed repair efficiency (27), base editing (28) and prime editing (29) will further increase the number and range of mutations that can be screened.

Our ability to perform GenIE ATAC experiments in a multiplex manner also allows us to understand more generally the functional consequences of variation within transcription factor recognition sequences in their endogenous context. This will allow us to interpret the effects of sequence or chromatin context on the effect of individual substitutions within transcription factor binding sites.

We thus believe that the GenIE-ATAC method will facilitate the prioritisation of candidate genetic variants from GWAS studies for further functional follow-up and provide mechanistic insights into how these polymorphisms affect gene expression and influence disease.

## Materials and Methods

### Experimental design

The GenIE-ATAC method requires the SNP of interest to be located within an ATAC peak in the assayed cell type. For each SNP, three primers were designed: an upstream biotinylated forward primer for the linear PCR enrichment step, a nested forward primer for the ATAC PCR step, and a reverse primer for gDNA amplification of the region (Fig S1). All primers bound within the ATAC peak, the forward primers did not overlap with each other and were located close to the SNP but outside the HDR template for editing experiments. The nested forward primer and reverse primer contained adaptor sequence for the addition of barcodes for Miseq (30) and amplicons were less than 295 bp to allow sequencing using a 150 PE Miseq run. All primers were unique in BLAT searches. For reverse experiments, equivalent primers were designed but in the opposite directions. For editing experiments, we chose the guide with a cut site closest to the SNP of interest and off-target cutting of guides was checked using WGE (https://www.sanger.ac.uk/htgt/wge/). All guide sequences, HDR oligo sequences, and primer sequences used are detailed in Supplementary Table 1 and 2.

### Tn5 production

GenIE-ATAC requires the Tn5 protein to be loaded with a single oligonucleotide (MEDS-A) which differs from the commercially produced Tn5 which is loaded with MEDS-A and B oligonucleotides. We purified Tn5 using a C-terminal intein tag and a chitin-binding domain as previously published (31), with some adaptations including the use of chitin magnetic beads. We transformed pTBX1-Tn5 plasmid (Addgene 60240) into C3013 cells (NEB) and expressed Tn5 as previously described. Bacterial pellets (from 500 ml culture) were resuspended in 12 ml cold HEGX (20 mM HEPES-KOH at pH 7.2, 0.8 M NaCl, 1 mM EDTA, 10% glycerol, 0.2% Triton X-100) plus proteinase inhibitors (PI) (Complete, Roche), sonicated on ice at 80% power for 8 x 30 s on/ 30 s off with a probe sonicator and centrifuged (13,000 rpm for 20 min 4°C) to produce 12.5 ml cleared lysate. Neutralised PEI (1 ml 10%, final concentration 0.8%) was added dropwise to the lysate to precipitate the DNA, and the lysate was centrifuged to clear (13,000 rpm for 20 min 4°C). The supernatant (~12 ml) was added to 1 ml HEGX washed chitin magnetic resin (NEB) and left to bind for 1 h at 4°C. Beads were washed 4 times with HEGX buffer before adding 250 nmoles (50 μl) of annealed MEDS-A oligo in 3 ml HEGX buffer plus PI and rocked overnight at RT.

#### Annealed MEDS

MEDS-A: TCGTCGGCAGCGTCAGATGTGTATAAGAGACAG

MEDS-REV: pCTGTCTCTTATACACATCT/intT

Oligos were resuspended in 10 mM Tris-HCl (pH 8.5) to a concentration of 10 nmoles/μl and 25 μl of each were mixed, heated to 95°C and left to cool slowly to RT. Importantly, annealed oligos were used on the same day.

After incubation with oligos, the beads were washed 4 times with HEGX buffer, and Tn5 protein was released from the beads by incubation with 50 mM fresh DTT in 4 ml HEGX for 48 h at 4°C. Eluted protein was collected and dialysed overnight in 2L 2x Tn5 dialysis buffer (100 HEPES-KOH at pH 7.2, 0.2 M NaCl, 0.2 mM EDTA, 2 mM DTT, 0.2% Triton X-100, 20% glycerol) at 4°C. The Tn5 was concentrated using 10K MWCO spin protein concentrator (Pierce) to 24 uM (1.32 mg/μl) as determined by Bradford. Then an equal volume of 100% glycerol was added and carefully mixed before aliquoting and storing at −20°C. Tn5 activity was titrated using gDNA in tagmentation buffer (5x buffer: 50 mM TAPS-NaOH, 25 mM MgCl_2_, 50% v/v DMF pH 8.5). The amount of Tn5 that was needed to tagment 1 μg gDNA to around 400 bp was found to be equivalent to the amount of Tn5 needed to be added to 0.5 x 10^6^ nuclei to tagment to the optimal size of fragments (around 400-800 bp) for GenIE-ATAC. This amount was between 1-5 μl of Tn5 produced as described above.

### hiPSC cell culture and ethics

iPSC lines were generated as part of the HipSci project (KOLF_2, Cambridgeshire 1 NRES REC Reference 09/H0304/77) and work on these is covered under HMDMC 14/013. Human KOLF_2 (HIPSCI, www.hipsci.org) were grown in feeder-free conditions in TeSR-E8 medium (StemCell Technologies) on Synthemax (Corning) (final amount 2 μg/cm^2^) and routinely passaged 1:10 every 5 days using Gentle Cell Dissociation Reagent (Stemcell Technologies).

### Arrayed CRISPR-Cas9 editing

hiPSCs were edited by nucleofection of ribonucleoprotein (RNP) complex containing full-length chemically modified synthetic guide RNA and SpCas9, along with a ssODN repair template (19, 21). Briefly, SpCas9 was expressed and purified from *E.coli* using a His-tag. Diluted SpCas9 (5 μl, 20 μg) was mixed with full-length guide RNA (Synthego) (5 μl, 225 pmoles) at RT for 20 min for RNP complexes to form, followed by addition of the ssODN repair template (5 μl, 500 pmoles) just before the nucleofection. Cells were dissociated using accutase and 1 x 10^6^ cells in P3 buffer were mixed with the RNP/template complex and nucleofected in large cuvettes (V4XP-3024 Lonza) using 4D-Nucleofector on program CA137. After nucleofection cells were plated onto a 10 cm dishes coated with Synthemax (5 μg/cm^2^) with TeSR-E8 supplemented with Rock inhibitor. After 24 hours, the media was exchanged for TeSR-E8 and after ~10 days the cells, when they had grown to approximately 80% confluence, were harvested by accutase and processed for ATAC (0.5×10^6^ cells) or pellets (2 x10^6^ cells) were frozen for gDNA extraction. Routinely at least 3 ATAC samples and 3 gDNA samples were analysed per pool of cells.

### ATAC assay, Linear PCR and nested PCR

hiPSCs grown on Synthemax were washed with PBS and 6 ml accutase was added for 5 min at 37°C. The accutase was aspirated and the cells were gently resuspended in 10 ml TeSR-E8. Cells were pelleted (300 g 3 min) and resuspended in 1 ml cold PBS. After counting, 0.5×10^6^ cells were transferred to an eppendorf (not loBind) and pelleted (300 g 3 min). The cells were then resuspended in 500 μl fresh ice cold sucrose buffer (10 mM Tris-HCl pH 7.5, 3 mM CaCl_2_, 2 mM MgCl_2_, 0.32 M sucrose) and incubated on ice for 12 min. Triton-X-100 was added to a final concentration of 0.5% (25 μl of a fresh 10% solution), mixed gently and incubated on ice for 6 min. The whole lysis solution was transferred to a new eppendorf (not loBind) and spun 450g for 5 min at 4°C to collect the nuclei. All traces of the lysis buffer were removed from the nuclei and 50 μl tagmentation mixture (10 μl 5x tagmentation buffer, Tn5 (amount from titration above), water to 50 μl) was added to the nuclei, gently resuspended and transferred to a Lobind eppendorf. Nuclei were tagmented for 30 min at 37°C and the DNA was cleaned up using a MinElute PCR cleanup kit (Qiagen) and eluted in 21 μl EB. The DNA was quantified using Qubit and was typically 30-60 ng/μl. For each SNP, 3-4 independent tagmentations were carried out from the same pool of cells.

Linear PCR was carried out on the purified DNA for each ATAC repeat:

|  |  |
| --- | --- |
| purified tagmented DNA | 10 µl |
| biotinylated forward primer (2uM) fresh | 1.5 µl |
| KAPA hotstart ready mix | 25 µl |
| Water | 13.5 µl |

1. 95°C 5 min
2. 98°C 20 s
3. 60°C 30 s
4. 72°C 30 s
5. Goto Step 2 49x

Kapa (non-hotstart) enzyme (1 μl) was spiked into the PCR mixtures and a further 50 cycles completed.

1. 98°C 20 s
2. 60°C 30 s
3. 72°C 30 s
4. Goto Step 1 49x
5. 72°C 10 min
6. 4°C storage

Biotinylated DNA pulldown was performed using Dynabeads MyOne Streptavidin C1 and 20 μl slurry was used per PCR. The beads were washed twice in 40 μl PBS with 0.1% BSA and once with 2xBW buffer (10 mM Tris-HCl, pH 7.5, 1 mM EDTA, 2 mM NaCl) before being resuspended in 50 μl 2xBW buffer. The whole linear PCR (50 μl) was added to the bead suspension and rotated overnight at RT. The beads were collected on the magnet, and the supernatant removed. The beads were washed once with 100 μl water, and resuspended in 30 μl EB.

The subsequent amplification of DNA used the beads as template with the nested forward primer specific for the SNP of interest and the Tn5A primer (TCGTCGGCAGCGTCAG) specific for the oligonucleotide loaded into the Tn5 and integrated into the DNA during tagmentation. The amount of beads was tested (Fig S4) and routinely we use 10 μl allowing up to 3 technical repeats to be performed per ATAC. PCR was carried out using PowerUp SYBR Green Master Mix (Applied Biosystems) with 10 μl bead template and 0.4 μM final concentration forward and reverse primers in a 50 μl reaction.

1. 95°C 10 min
2. 95°C 15 s
3. 57°C 15 s
4. 72°C 30 s
5. Goto Step 2 35x
6. 72°C 5 min
7. 4°C storage

The PCR products were analysed on a 2% agarose gel, stained with ethidium bromide, and should be a smear of DNA sizes ranging from ~400-800 bp (similar to the tagmented DNA).

### gDNA extraction and PCR

gDNA was prepared using the MagAttract HWM kit (Qiagen) (19). PCR was carried out using PowerUp SYBR Green Master Mix (Applied Biosystems) with 5 μl gDNA (250 ng - 500 ng) template and 0.4 μM final concentration forward and reverse primers in a 50 μl reaction. Typically 3-4 PCRs were carried out from one gDNA preparation. We also tested using tagmented gDNA as the template for PCR (Fig S2) and in these experiments 250-500 ng DNA was tagmented as described in the ATAC experiment, cleaned up using the MinElute PCR cleanup kit (Qiagen) and used as PCR template.

### Barcoding and sequencing

In order to add the Illumina indices (30) we performed a second PCR using 1 μl PCR1 (from ATAC or gDNA samples), PowerUp SYBR Green Master Mix (Applied Biosystems), and 0.4 μM final concentration forward and reverse primers in a 25 μl reaction.

### CTCF saturation mutagenesis experiment

To carry out the editing for the saturation mutagenesis of CTCF binding site, we mixed 9 HDR oligos equally and performed 8 nucleofections with RNP as described above. The cells were pooled and grown over 5 x 10 cm dishes and 9 ATAC and 9 gDNA independent samples were harvested. For the linear PCR we inputted 20 μl tagmented DNA, and after biotin pulldown, we used all 30 μl beads into the PCR (split over 4 PCR reactions). These PCRs were pooled before adding the Miseq barcodes. For the gDNA PCRs, we inputted 4x more gDNA and split over 4 PCR reactions before pooling for Miseq barcoding. This was to make sure we had good coverage of all of the editing events that had taken place in the pool of cells.

### Sequencing and read alignment

PCR amplicons were sequenced using 2 x 150 bp reads on an Illumina MiSeq instrument. Since amplicons were all smaller than 295 bp, we merged the overlapping 150-bp paired-end reads using FLASH v1.2.11 (32) to improve alignment of Cas9-induced deletions. As input to FLASH we specified a minimum overlap of 10 bp, fragment size as the amplicon size, fragment standard deviation of 40, and maximum mismatch density of 10%, and used the --allow-outies parameter. A mean of 98.6% of reads could be successfully merged, with standard deviation of 1.5%. For all regions except rs141252451 in MIR8070, we aligned merged reads to the GRCh38 human reference sequence using bwa mem v0.7.17 (33), with lenient parameters to allow aligning Cas9-induced deletions (-O 24,48 −E 1 −A 4 −B 16 −T 70 −k 19 −w 200 −d 600 −L 20 −U 40). For rs141252451, we aligned to an amplicon sequence that included the 2-bp insertion defined by rs141252451, since rgenie excludes insertion reads by default, assuming these reflect abnormal events for Cas9-mediated editing.

### Analysis with rgenie

The core analysis performed in rgenie is the same as previously described (19). Briefly, for each replicate (cDNA or gDNA) at each locus, rgenie extracts reads mapping to the targeted region from the aligned BAM file. To quantify the effect of an allele X on the ATAC-seq readout, we first determine for each replicate the ratio of the read count of X to that of the WT allele, r = (read count X / read count WT). We separately compute the mean of this ratio for ATAC replicates, r^A^, and for gDNA replicates, r^G^. If allele X alters chromatin accessibility, then the ratios will differ between ATAC and gDNA, i.e. r^A^ ≠ r^G^. We use a two-tailed unequal variances t-test to test for a difference in this ratio, and we report p-values from this test. Although the statistical analysis is identical for an RNA or an ATAC readout, we updated rgenie to provide plots and statistical summaries tailored to an ATAC analysis. Specifically, an ATAC analysis by default reports only deletions wholly contained within a +/− 6 bp window around the targeted SNP. This is because with ATAC-seq the reads span a range of sizes depending upon the ATAC peak shape, with most reads tending to be short. As a result, longer deletions are not reliably captured; this differs from RNA, where all reads span the full amplicon. We also highlight allele effect statistics for the two most prevalent deletions within this window. In Supplementary Tables 3 and 4, we provide a statistical analysis summary from applying rgenie to the 8 SNPs and the CTCF mutagenesis experiments, respectively. See the Data Availability section for links to rgenie input file details, amplicon sequences, and HDR/WT alleles for these analyses.

## Acknowledgements

pTXB1-Tn5 was a gift from Rickard Sandberg (Addgene plasmid # 60240 ; http://n2t.net/addgene:60240 ; RRID:Addgene_60240). We acknowledge Scientific Operations at the Sanger Institute for support in next generation sequencing and quality control. We thank members of the Bassett lab for helpful discussions.

## Data Availability

The R package (rgenie) developed in this study is available at https://github.com/Jeremy37/rgenie Raw data is available at Zenodo (https://doi.org/10.5281/zenodo.5802511).

## Authors’ contributions

S.C. and J.S. contributed equally to this work. S.C. planned and conducted experiments with help from E.C. and Q.W.. J.S. developed the software and conducted the analyses. A.B. supervised the study.

