## Supplementary Figures for "Screening for functional regulatory variants in open chromatin using GenIE-ATAC"

### Figure S1

#### ATAC samples

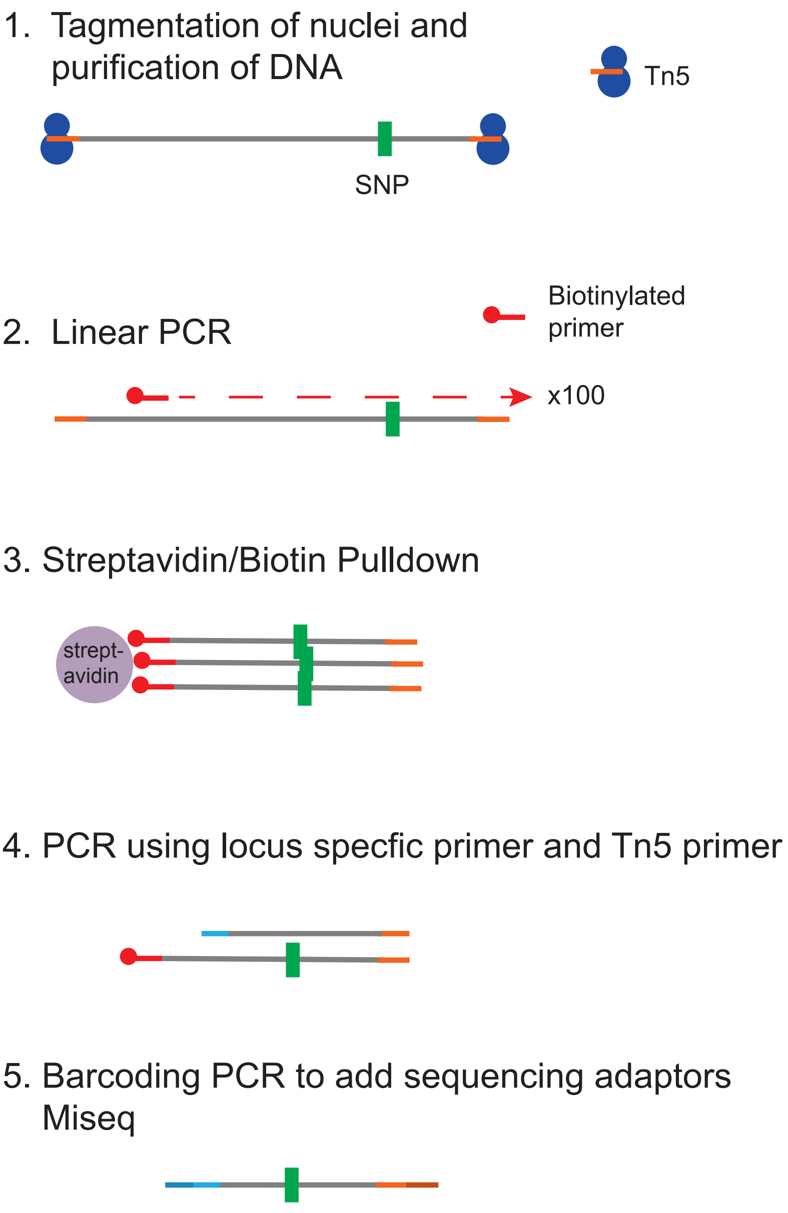


**Figure S1. Schematic of ATAC amplicon sequencing.** After tagmentation, a linear PCR is carried out using a biotinylated primer to allow enrichment of the DNA surrounding the SNP of interest using a biotin pulldown. The locus specific amplicon is then amplified by PCR using the common Tn5 MEDS-A primer and a gene specific nested primer, before adding Miseq adaptors for sequencing.

#### Figure S2

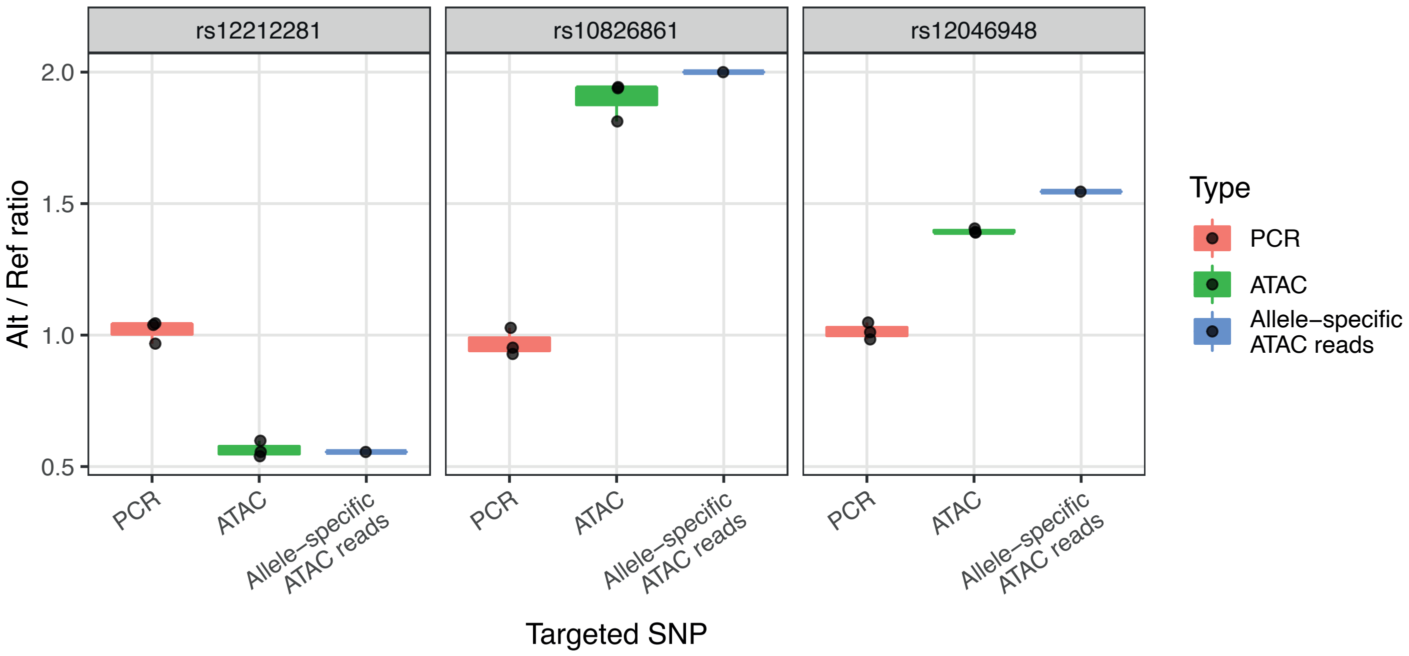


**Figure S2. GenIE-ATAC produces identical results when linear PCR is performed in the opposite direction.** GenIE-ATAC was also performed on the three positive control SNPs shown in Fig 2 in KOLF_2 hiPSCs, with the primers (linear PCR and nested) designed in the opposite orientation, with the Tn5 being integrated on the other side of the SNP of interest. Graphs show the ratio of chromatin accessibility of the alt allele normalised to the reference genome sequence (Alt/Ref ratio). Red bars indicate amplicon sequencing of gDNA (PCR), green bars from amplicon sequencing of ATAC material (ATAC), blue bars show allele-specific reads from ATAC-seq data (allele-specific ATAC reads) from KOLF_2 iPSCs. Individual repeats are indicated with dots; boxplot hinges represent the 25th and 75th percentiles, where these are interpolated due to the small number of points (n=3), and whiskers extend to the most extreme data point not further than 1.5 times the inter-quartile range from the hinge.

#### Figure S3

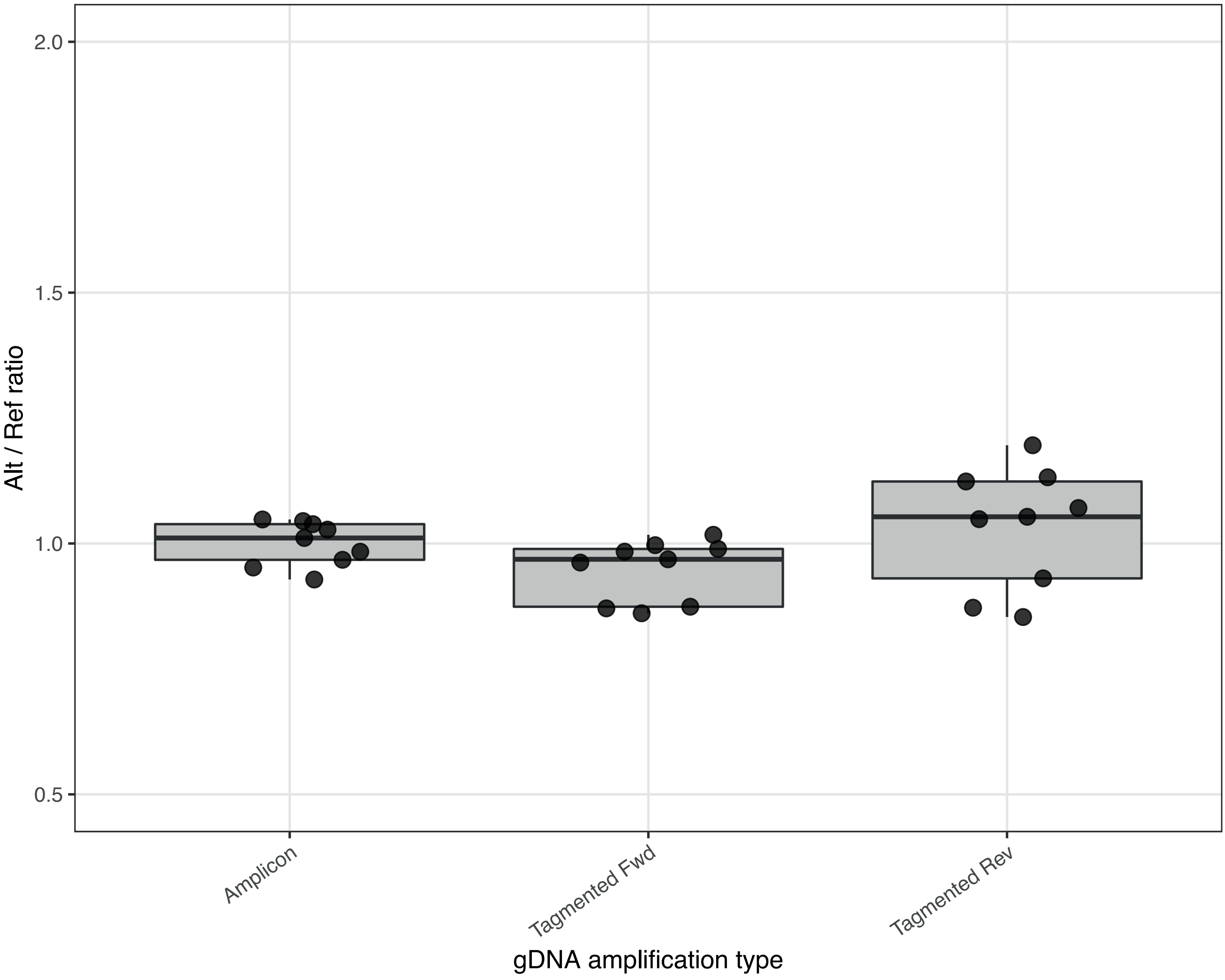


**Figure S3. Tagmented gDNA increases variation compared to untagmented gDNA.** In the heterozygote experiment shown in Figure 2 where the ratio of Alt/Ref reads would be expected to be 1, amplicon sequencing was carried out using either gDNA (amplicon), tagmented DNA in forward direction (tagmented Fwd), or tagmented DNA in reverse direction (tagmented Rev, Fig S2) as template. Boxplot hinges represent the 25th and 75th percentiles of the alt/ref ratio, and whiskers extend to the most extreme data point not further than 1.5 times the inter-quartile range from the hinge.

#### Figure S4

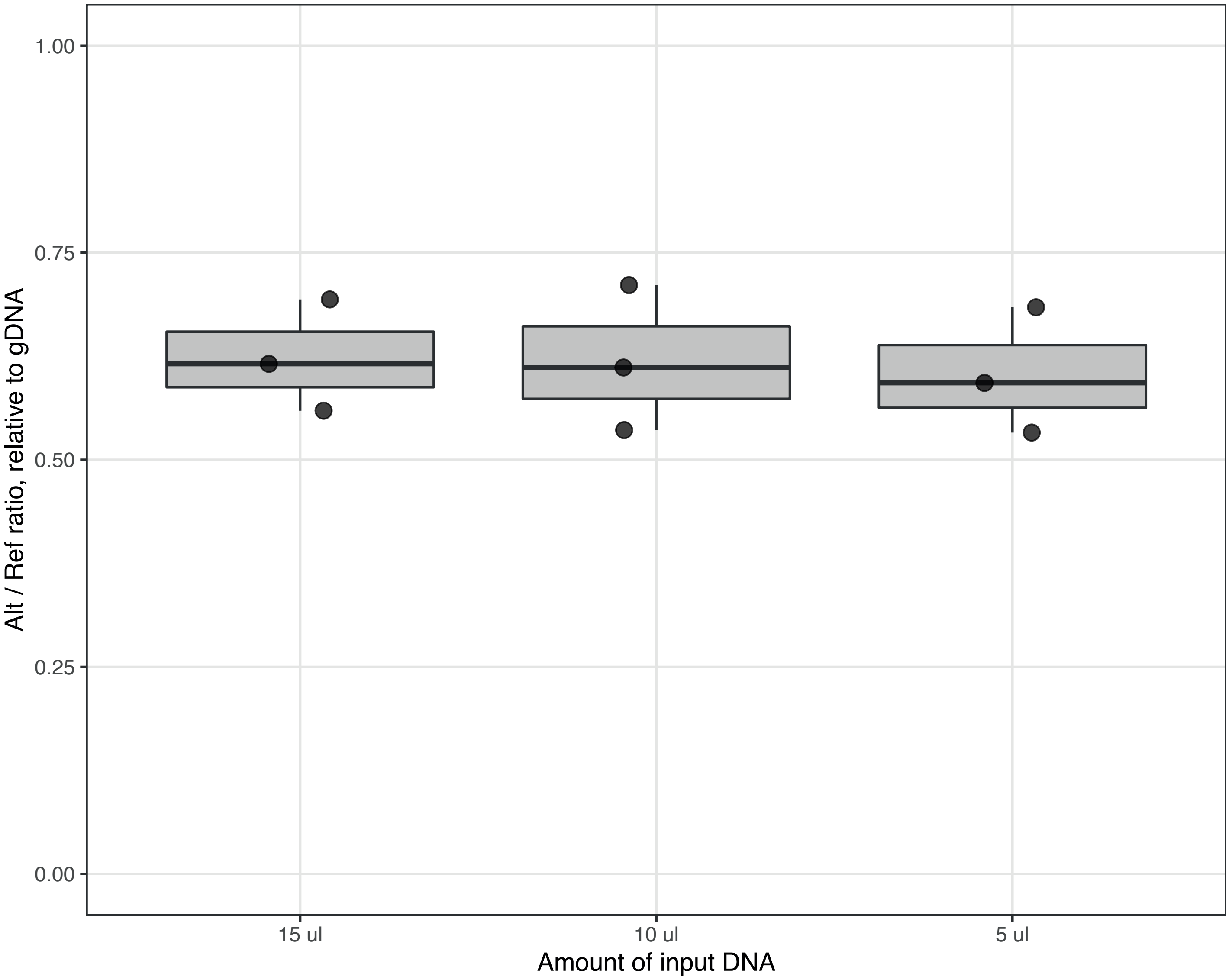


**Figure S4. Reducing the ATAC input DNA into the linear PCR does not increase variation.** In the editing experiment for SNP rs12269414 shown in Fig 3, differing amounts of ATAC DNA was added to the linear PCR reaction (15 µl, 10 µl or 5 µl). Boxplot hinges represent the 25th and 75th percentiles, where these are interpolated due to the small number of points (n=3), and whiskers extend to the most extreme data point not further than 1.5 times the inter-quartile range from the hinge.

**Figure S5**

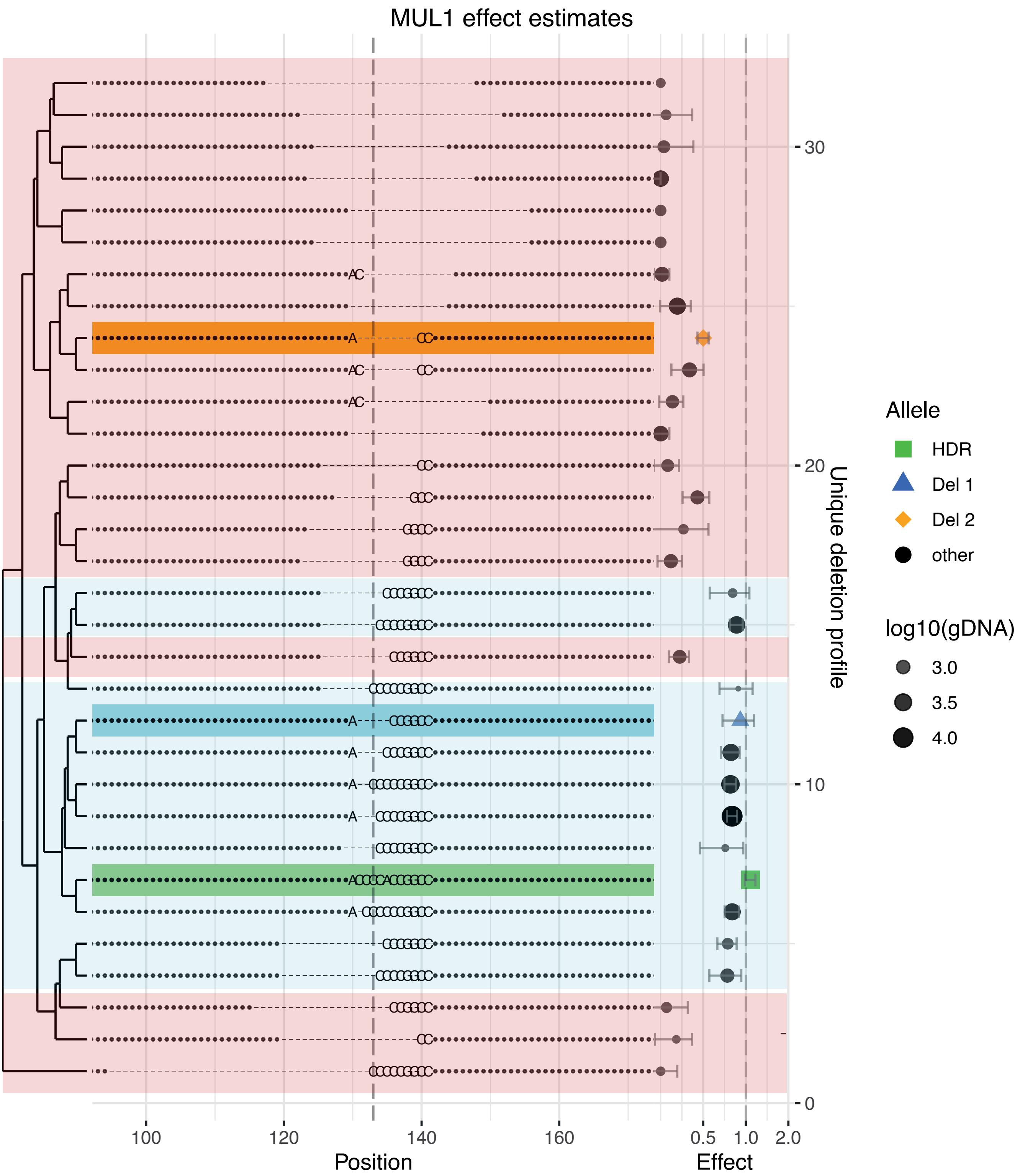


**Figure S5. Analysis of individual deletions around rs6700034 shows the importance of polyC tract in transcription factor binding.** Deletion profiles from the most prevalent alleles by read count from GenIE-ATAC targeting of rs6700034. The effect on chromatin accessibility for each allele relative to WT is indicated on the right. The HDR allele is highlighted in green. Del 1 and Del 2 (highlighted in blue and orange respectively) are small defined deletions around SNP of interest. Two classes of deletion alleles are highlighted: those in light blue have no effect change and keep transcription factor binding intact; those in pink show a strong reduction in chromatin accessibility and abolish transcription factor binding.
